# Multisensory gamma stimulation enhances adult neurogenesis and improves cognitive function in male mice with Down Syndrome

**DOI:** 10.1101/2024.10.03.616486

**Authors:** Md Rezaul Islam, Brennan Jackson, Maeesha Tasnim Naomi, Brooke Schatz, Noah Tan, Mitchell Murdock, Dong Shin Park, Daniela Rodrigues Amorim, Fred Jiang, S. Sebastian Pineda, Chinnakkaruppan Adaikkan, Vanesa Fernandez, Ute Geigenmuller, Rosalind Mott Firenze, Manolis Kellis, Edward S. Boyden, Li-Huei Tsai

**Affiliations:** Picower Institute for Learning and Memory, Massachusetts Institute of Technology, Cambridge, MA, USA; Department of Brain and Cognitive Sciences, Massachusetts Institute of Technology, Cambridge, MA, USA; Institute for Medical Engineering and Science, Massachusetts Institute of Technology, Cambridge, MA, USA; Departments of Biological Engineering and Brain and Cognitive Sciences, McGovern Institute, Cambridge, MA, USA; MIT Computer Science and Artificial Intelligence Laboratory, Cambridge, MA, USA; Broad Institute of MIT and Harvard, Cambridge, MA, USA; Koch Institute, Massachusetts Institute of Technology, Cambridge, MA, USA; Howard Hughes Medical Institute, Massachusetts Institute of Technology, Cambridge, MA, USA

**Keywords:** Multisensory gamma stimulation, 40 Hz, single nuclei RNA-seq, gene-regulatory network, Down syndrome, non-invasive therapy

## Abstract

Down syndrome (DS) has been linked with deficits in hippocampal dependent cognitive tasks and adult neurogenesis, yet treatment options are still very limited. We and others previously showed that a non-invasive multisensory gamma stimulation using light and sound at 40 Hz ameliorated Alzheimer’s disease pathology and symptoms in mouse models. In this study, we tested the effects of 40 Hz multisensory stimulation in the Ts65Dn mice, a mouse model of DS. For three weeks, mice were exposed daily to one hour of stimulation or one hour of ambient light and sound. Mice receiving the stimulation showed improved object recognition and spatial working memory. Using single nuclei RNA-seq and experimental validations in mouse hippocampal samples, we identified underlying expression changes in gene regulatory networks and demonstrated increased adult neurogenesis and reorganization of synapses as potential mechanisms for these improved cognitive phenotypes. Together, our data reveal a novel effect of multisensory gamma stimulation on adult neurogenesis and beneficial effects of 40 Hz treatment on cognitive function in DS model mice.

**Significance Statement:** We present strong evidence, using a well-characterized mouse model, that the cognitive and neurogenesis deficits in Down syndrome can be improved through non-invasive multi-sensory gamma stimulation. Employing a systems biology approach, we provide extensive hippocampal single-cell resolution gene expression signatures and changes in gene regulatory networks in response to sensory gamma stimulation.

## Main Text

### Introduction

Down syndrome (DS) has been linked with impaired neurogenesis [1] and aberrant synaptic functions in the hippocampus [2]. These changes are consistent with the reported deficits in memory and other cognitive tasks in individuals with DS [3,4]. Additionally, a recent study found dementia to be associated with mortality in 70% of older adults with DS [5]. However, there are currently limited therapeutic options available to manage cognitive performance in individuals with DS. We have previously reported the utility of using sensory stimulation presented at gamma frequency for combating cognitive decline in mouse and human subjects [6–8]. The sensory gamma stimulation reduced Alzheimer’s disease (AD) related pathology, improved neuronal survival and synaptic density, and enhanced cognitive function in multiple mouse models of AD [7,9,10]. In a more recent literature, we showed that the reduced pathology was specifically observed at 40 Hz stimulation but not at 8 or 80 Hz stimulation [9]. Others have also reported the beneficial effects associated with gamma stimulation in AD [11–13] as well as in other neurological diseases such as Parkinsońs disease [14]. However, the efficacy of sensory gamma stimulation at 40Hz has not been tested in DS. The Ts65Dn mouse, the most widely used model of DS, contains an extra chromosome spanning most of the distal region of mouse chromosome 16 homologous to human chromosome 21 [15]. This murine model recapitulates phenotypes present in individuals with DS including synaptic abnormalities [16] and deficits in hippocampal-dependent spatial learning and memory [15]. As such, this murine model provides invaluable opportunities for exploring and testing potential therapeutic interventions for cognitive deficits in DS. Here, we used this well-characterized DS mouse model to test the effects of sensory gamma stimulation on cognitive benefits within the DS model and observed that this treatment improved cognitive function, enhanced adult neurogenesis, and induced expression changes in genes involved in synaptic organization.

### Results

#### Multisensory gamma stimulation improves cognitive performance of Ts65Dn mice

To examine the effect of the sensory gamma stimulation on cognitive performance in DS, we used the Ts65Dn mouse model of DS, which recapitulates phenotypes present in individuals with DS including synaptic abnormalities [16] and deficits in hippocampal-dependent spatial learning and memory [15] and adult neurogenesis [17]. We exposed one group of Ts65Dn mice to one hour of 40 Hz auditory and visual stimulation per day (stimulation), while a parallel group was exposed to ambient light and sound (ambient, **Fig 1A**). Littermate wild-type controls were not included in this experimental setup since the well-established cognitive deficits of Ts65Dn mice in hippocampus-dependent tasks, as reported in previous studies [15,18,19], offer a clear benchmark for evaluating the effects of stimulation on cognitive improvements. Shortly before the completion of the three-week treatment period using the 40 Hz or ambient auditory and visual stimulation, we conducted the Novel Object Recognition (NOR), Novel Object Location (NOL), and elevated Y-maze behavioral tests to evaluate the effects of the stimulation on object recognition (**Fig 1B**) and spatial memory (**Fig 1C-D**). For the NOR/NOL tests, both groups of mice were first presented with two objects in fixed locations. When one of the familiar objects was later replaced by a novel object in the same location (NOR, **Fig 1B**) or placed in a novel location (NOL, **Fig 1C**), the stimulation group spent a significantly higher percentage of time than the ambient group exploring the novel object (stimulation = 64.1+/-12.3%, ambient = 48.6+/-21.5%, p = 0.04) and the familiar object in the novel location (stimulation = 73.4+/-11.7% vs ambient = 46.6+/-15.4%, p < 0.001). Importantly, the observed changes in performance between stimulation and ambient groups were not related to differences in locomotory behaviors in an open field area (**Fig S1A, Fig S1B**). Thus, the NOR/NOL results indicate a clear improvement in both object recognition and spatial memory in the stimulation group compared to the ambient group. Spatial memory was further assessed by spontaneous alternation in the elevated Y-maze test, which allows mice to freely explore the three arms of a Y-shaped maze. Consecutive entries into three different arms are considered a spontaneous alternation, expressed as % of all possible entry triads. Mice with stimulation demonstrated a significantly higher number of spontaneous alternations than the ambient group (stimulation = 61.15+/-9.1%, ambient = 51.3+/-8.7%, p = 0.02, **Fig 1D**), while the number of total entries was not significantly different between the two groups (stimulation = 29.2+/-7.9, ambient = 24.1+/-6.3, **Fig 1D**). These results indicate improved spatial working memory in DS mice in response to stimulation. Since both object recognition and spatial memory are hippocampus-dependent memory tasks, the behavioral testing results also imply that the sensory stimulation impacts hippocampal neurons. This was experimentally confirmed by performing staining for c-Fos, a widely used marker of neuronal activity [20], on brain sections from mice in the stimulation and ambient groups. Mice in the stimulation group showed an increased proportion of c-Fos positive nuclei in the hippocampus compared to mice in the ambient group (**Fig S2**).

**Fig 1.**
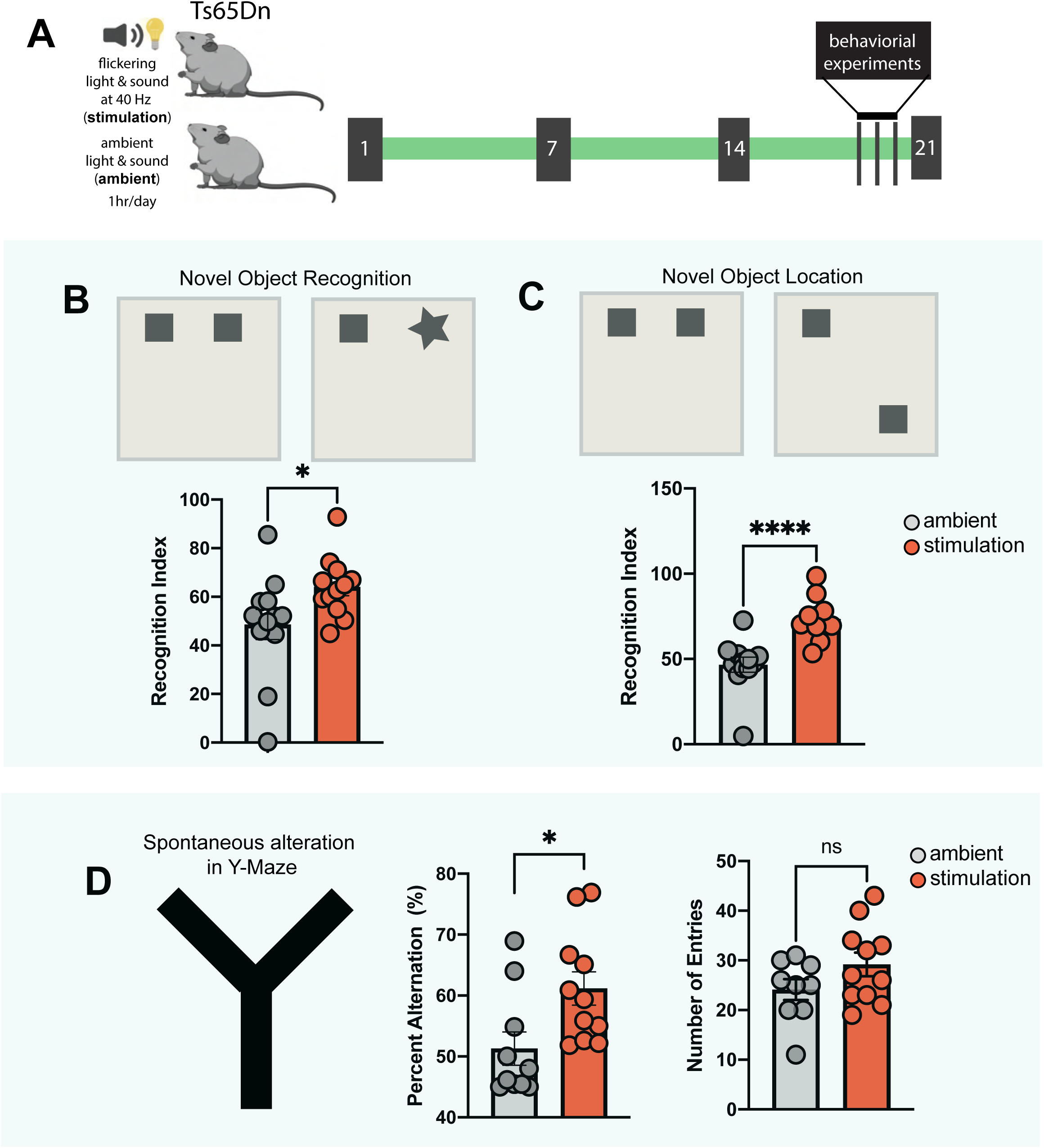
Multisensory gamma stimulation improves spatial working memory in Ts65Dn mice. A) Experimental scheme. B) Novel Object Recognition (NOR) test. Quantification of recognition index during NOR. Average Novel object exploration time: ambient: 9.73 seconds, stimulation: 12.84 seconds. Average familiar object exploration time: ambient: 10.31 seconds, stimulation: 7.20 seconds. C) Novel Object Location (NOL) test. Quantification of recognition index during NOL. Average Novel location exploration time: ambient: 9.33 seconds, stimulation: 14.72 seconds. Average old location exploration time: ambient: 10.70 seconds, stimulation: 5.34 seconds. D) Spontaneous alteration based on Y-maze. Quantification of percent alteration and total number of entries. ambient = 12, stimulation = 12, unpaired t-test, two-tailed, *P<0.05, ****P<0.0001. Error bars indicate mean ± standard error mean.

#### Multisensory gamma stimulation increases expression of synapse-related genes in excitatory neurons in the hippocampus of Ts65Dn mice

To decipher potential mechanisms underlying sensory stimulation-mediated cognitive improvement in Ts65Dn mice, we performed single-nucleus RNA sequencing (snRNA-seq) on hippocampal samples from the stimulated and ambient groups of mice (**Fig 2A)**, yielding 15,884 nuclei with high quality transcriptomic profiles (**Fig S3**). Based on the top variable genes, the nuclei were categorized into transcriptionally distinct clusters that could be visualized in uniform manifold approximation and projection (UMAP) space and assigned to principal brain cell types based on the expression of canonical markers (**Fig 2B, Fig S3E**). The majority (69.7%) of captured nuclei represented neurons (54.5% excitatory and 15.2% inhibitory), 26.5% were derived from glial cells, and the remaining 3.8% represented vascular and choroid plexus cells (**Fig S3F**). Samples were well integrated across both groups (**Fig S3G**), and broad cell type constituency did not differ significantly between the stimulation and ambient groups (**Fig S3H)**.

**Fig 2.**
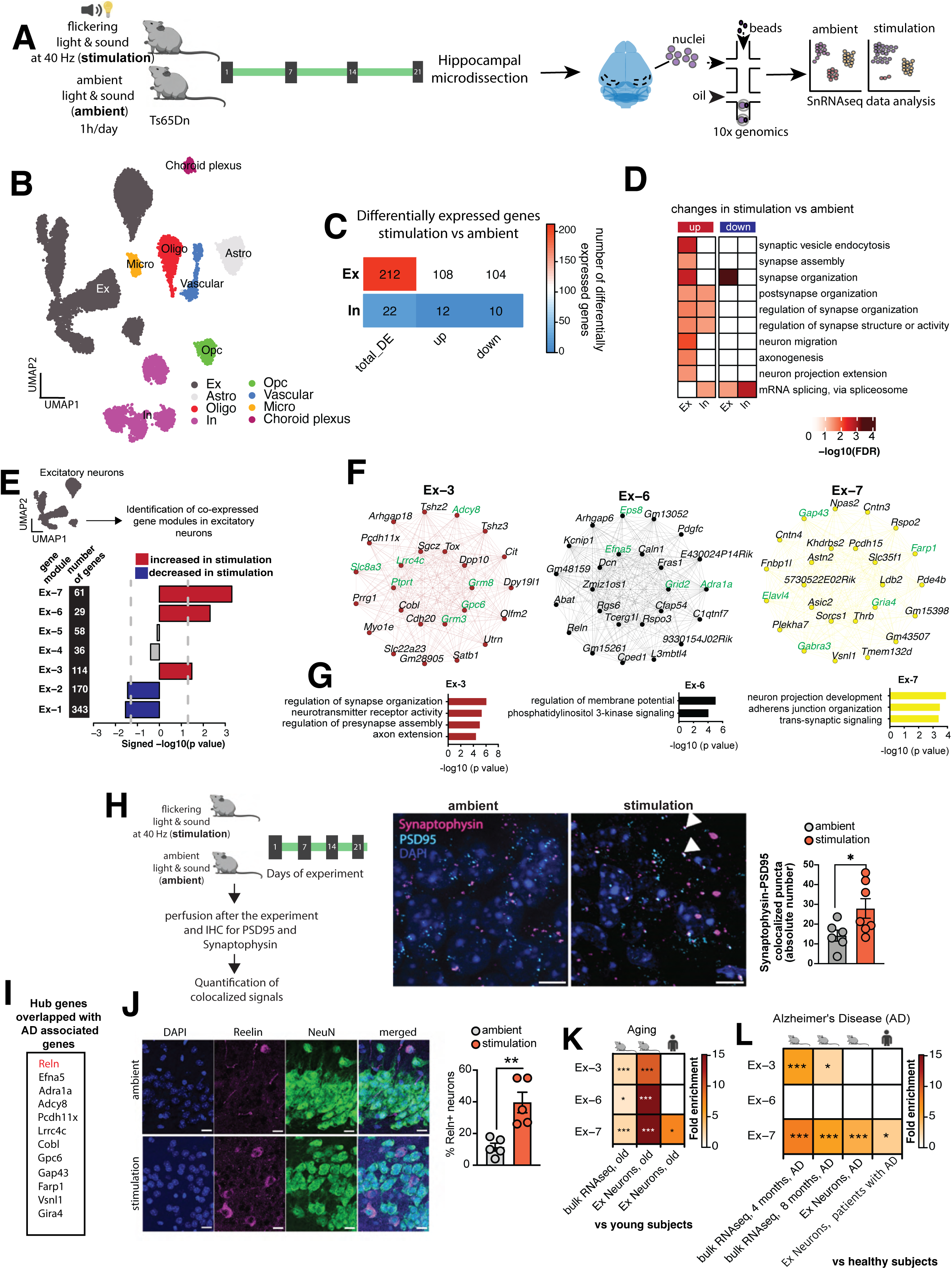
Molecular signatures underlying multisensory gamma stimulation-associated cognitive benefits in the hippocampus. A) Experimental scheme for the single nuclei RNA-seq experiment in Ts65Dn mice B) Unbiased clustering of hippocampal snRNA-seq data for 15884 nuclei represented on a UMAP. Cell types are annotated based on the marker genes. Ex: Excitatory Neurons, Astro: Astrocytes, Oligo: Oligodendrocytes, In: Inhibitory Neurons, OPC: Oligodendrocyte precursor cells, Micro: Microglia. C) Heatmap showing number of differentially expressed genes in excitatory and inhibitory neurons in the hippocampal snRNA-seq data from stimulation vs ambient Ts65Dn mice. D) Gene ontology analyses for the upregulated and downregulated genes in hippocampal excitatory and inhibitory neurons in stimulation vs ambient Ts65Dn mice. Color represents statistical significance after multiple testing adjustments. E) Signed bar plot displaying significant co-expression changes of the identified 7 gene modules in hippocampal excitatory neurons in stimulation vs ambient TsD65n mice. Red color denotes increased and blue color decreased expression in stimulated mice. X-axis represents signed -log10 p value. F) Hub genes of modules Ex-3, Ex-6 and Ex-7. Genes that encode synaptic proteins are highlighted in green. G) Top gene ontology biological processes for modules Ex-3, Ex-6, and Ex-7. H) Quantification of mature synapses in granule cell molecular layer of dentate gyrus in stimulation and ambient Ts65Dn mice via co-localization of PSD95 and synaptophysin. Representative images of mouse brain slices stained with PSD95, synaptophysin and DAPI. Images were acquired using 63x objective. Triangles indicate colocalized puncta. Y-axis represents the absolute number of colocalized puncta. Scale bar = 10 microns. t-test, two-tailed, unpaired, *P<0.05. Error bars indicate mean ± sem. ambient = 6, stimulation = 7 mice. I) Hub genes overlapping with disease risk genes for Alzheimeŕs disease. J) Quantification of *Reln*+ neurons after ambient or 40Hz stimulation. Representative confocal images of hippocampal CA3 area brain slices from Ts65Dn mice after staining for Reelin and NeuN with DAPI counterstain. Merged panel shows all three markers. 20x objective. Scale bar = 50 pixels. Bar plots show the mean proportion of *Reln*+ neurons between the ambient and stimulation groups. Error bars indicate mean ± sem. t-test, two-tailed, unpaired, **P<0.01. Number of mice = 5/group. K-L) Hypergeometric overlap of the genes in modules Ex-3, Ex-6, and Ex-7 with different sets of downregulated genes related to (K) aging and (L) Alzheimer’s disease (AD). Color represents fold enrichment. *FDR<0.05, **FDR<0.01, ***FDR<0.001.

We then performed differential gene expression analyses focusing on the neuronal clusters. Excitatory neurons showed a higher number of gene expression changes in response to the stimulation than inhibitory neurons, with 108 genes upregulated and 104 genes downregulated in the stimulation group compared to the ambient group (**Fig 2C**, **Fig S3I**, **Table S1**). Using a curated list of synaptic genes as reference [21], we found that synaptic genes were significantly overrepresented within the set of 108 upregulated genes (Fisher’s exact test, p value = 0.004; **Table S2**). Differentially upregulated synaptic genes included *Actb, Actg1, Atp6voc, Camk2b, Frrs1l, Gphn, Gpm6a, Hspa8, Il1rapl1, Mdga2, Nptn, Nrg1, Rab14, Rock1, Syt11*, and *Tuba1a*, of which several have been implicated in synaptic functions [22], synapse development [23], or synaptic plasticity [24]. Gene ontology (GO) analyses further demonstrated that the genes differentially expressed in excitatory neurons in response to stimulation were associated with various synaptic functions, with the majority related to synapse organization (e.g., synaptic vesicle endocytosis, synapse assembly, synapse organization, post-synapse organization, regulation of synapse organization, and regulation of synapse structure) (**Fig 2D**), and that some of these processes were also altered in inhibitory neurons (**Fig 2D**). Of note, reanalysis of published datasets (see Methods) unveiled downregulation of synapse organization associated genes in the brains of Ts65Dn mice (**Fig S4A-B, Table S3**) as well as in excitatory neurons in individuals with DS (**Fig S4C, Table S3**) compared to healthy controls.

To strengthen these findings in excitatory neurons and for additional insights, we employed an orthogonal computational approach. Using a weighted gene co-expression analysis (WGCNA), we identified seven gene regulatory modules in hippocampal excitatory neurons (**Fig 2E, Table S4, Fig S5A-B**). Three of these seven gene modules (Ex-3, Ex-6, Ex-7) showed significantly increased expression in the stimulation group compared to the ambient group, two (Ex-1 and Ex-2) displayed decreased expression (**Fig 2E**, **Fig S5C**), and two (Ex-4, Ex-5) showed no expression changes (**Fig 2E**). Focusing our downstream analyses on the three upregulated gene modules, we found that several of the hub genes encode synaptic proteins (**Fig 2F**) and few of these genes are implicated in synaptic organization (Farp1 [25]), synapse formation (Ptprt [26]) and synapse development (Lrrc4c [27], Efna5 [28]). Therefore, increased expression of these three gene modules may alter functional synapses in the hippocampus of the stimulation group.

To test this hypothesis experimentally, we performed immunostaining for the post-synaptic marker PSD95 and the pre-synaptic marker synaptophysin in the granule cell molecular layer of the dentate gyrus in the hippocampi from the ambient and stimulation groups (**Fig 2H**). Co- localization of both markers served as the readout of putative synapses. The stimulation group showed a significant increase in the number of such putative synapses in the dentate gyrus (**Fig 2H**), while no significant increase in the number of such synapses was seen in the CA1 region of the hippocampus (**Fig S6**). These findings confirm that the gene expression changes observed in response to stimulation lead to region-specific modification in synaptic organization within the hippocampus.

To better understand the molecular mechanisms, we focused on gene co-expression analyses rather than differential expression analyses for downstream analyses. Gene co-expression analyses enable the investigation of gene expression changes in a systems level framework [29]. Moreover, such analyses have potential to identify gene regulatory networks underlying complex biological phenotypes [30]. Additionally, they allow for comparative data analyses with existing datasets; and the hub genes identified based on co-expression analyses are potentially the driver of the gene circuits and provide critical insights into the underlying biological mechanisms [31,32]. Further analyses of the gene modules unveiled several hub genes that are associated with Alzheimeŕs Disease (AD) related dementia (**Fig 2I)**. Among those, *Reln* in particular has been implicated in AD [33,34] and aging [35], and a gain-of-function variant in this gene has been shown to confer resilience against autosomal dominant AD [36]. Remarkably, IHC unveiled an increased proportion of *Reln*+ neurons in the CA3 (**Fig 2J**) and DG (**Fig S7**) regions but not in the CA1 region (**Fig S7**) of the hippocampus of our stimulation group compared to the ambient group (**Fig 2J**).

Based on these observations and to further validate the significance of the three gene modules in cognitive benefits, we examined their expression patterns in published transcriptomic datasets (see Methods) focusing on aging and AD, both known risk factors for cognitive decline. Interestingly, we observed reverse expression pattern of these gene modules in context to the cognitive decline. Specifically, all three modules showed a steady decrease in expression with increasing age (**Fig S8**). In line with this, the gene modules particularly module Ex-3 and Ex-7 (both increased after stimulation), significantly overlapped with genes downregulated in excitatory neurons with aging (**Fig 2K, Table S5**) or AD (**Fig 2L, Table S6**) in mice and humans. Together, these data further confirm that gamma stimulation can increase expression of cognitively relevant gene modules in excitatory neurons and thus potentially contribute to the observed cognitive benefits.

#### Multisensory gamma stimulation increases adult neurogenesis in Ts65Dn mice

Since the Ex-7 (61 genes) module most consistently displayed overlap across aging and AD datasets in both mouse and humans (Fig 2K, 2L), we analyzed the transcription factors (TFs) for this module to identify upstream regulators. Among the TFs whose targets overlap with the module genes, TCF4 emerged as the highest ranked transcription factor when using a p value based on a hypergeometric test (**Fig 3A**, **Table S7**). Additionally, predicted TCF4 targets such as Nrg1, Rfx7, Robo2, Ttc3 were upregulated in excitatory neurons based on the differential expression analysis (**Table S1A**). This finding suggests that the differential gene expression and gene co-expression analyses converge on a common upstream gene regulatory factor. Interestingly, TCF4 is critical for progenitor proliferation [37], the peak of TCF4 expression coincides with neurogenesis [38], and it regulates neurogenesis and neuronal migration [39]. Moreover, TCF4 is a critical regulator that facilitates adult neurogenesis [40], and deficits in TCF4 expression lead to impaired adult hippocampal neurogenesis in mice [41]. Thus, we focused on TCF4 expression in the dentate gyrus of the hippocampus, a brain region that has been linked to adult neurogenesis. Excitingly, we observed dentate gyrus-specific increased TCF4 expression in the Ts65Dn mice after gamma stimulation (**Fig 3B, Fig S9**). Thus, we hypothesized that the stimulation might have an impact on adult neurogenesis.

**Fig 3.**
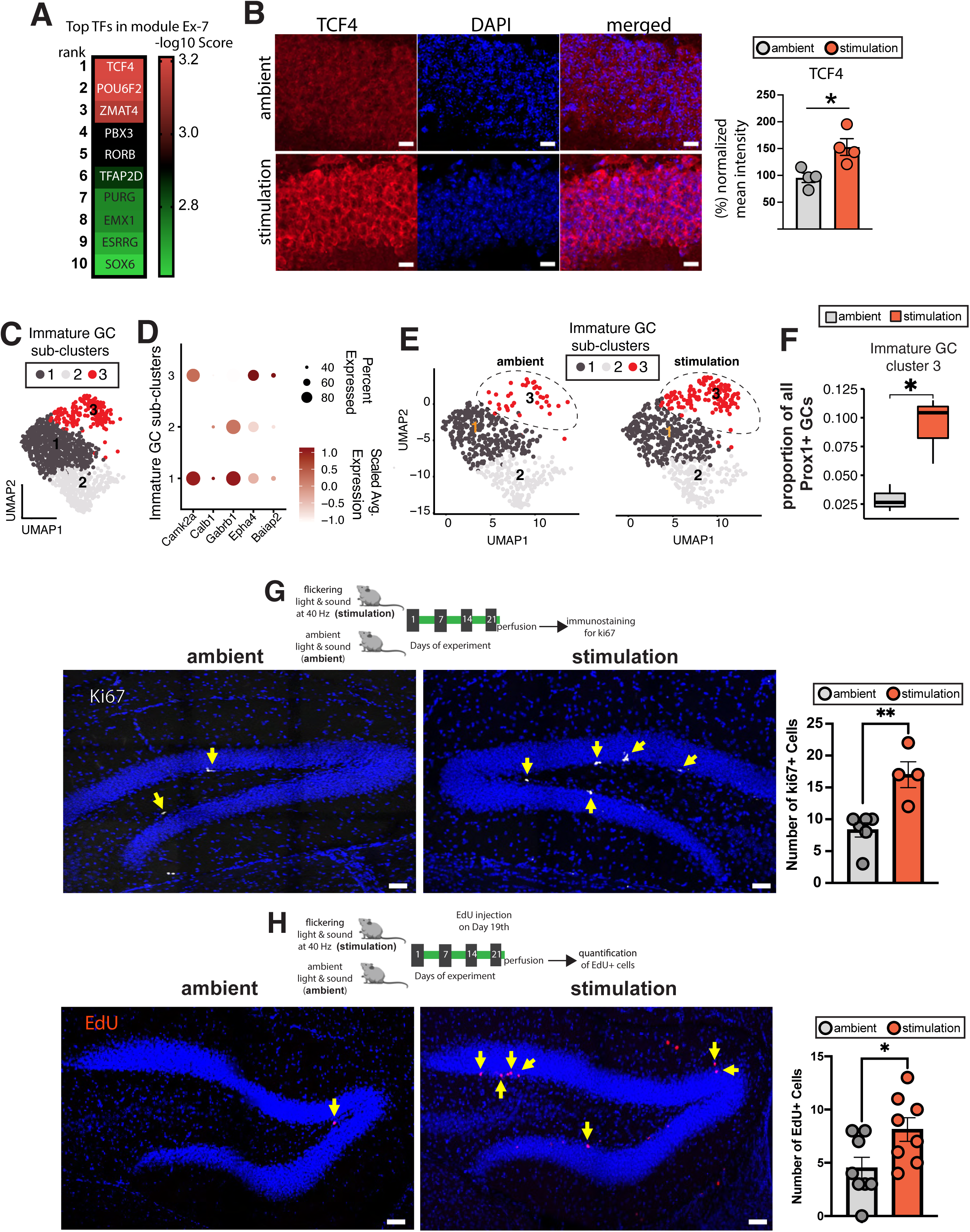
Multisensory gamma stimulation increases adult neurogenesis in Ts65Dn mice. A) Top 10 ranked TFs for module Ex-7. Color represents -log10 enrichment score based on ChEA3 analysis. B) Mean TCF4 fluorescence intensity after treatment of mice with or without 40 Hz stimulation. Representative images of TS65Dn mouse brain slices from the hippocampal granule cell molecular layer of dentate gyrus area after staining for TCF4 (green) and DAPI (blue). Merged panel shows both markers. Scale bar = 15 µm. Bar plots show normalized mean intensity (% of ambient group) between the two groups. t-test, two-tailed, unpaired. *P<0.05. number of mice = 4/group. C) Sub clusters of immature granule cells represented on a UMAP D) Dot plots showing relative enrichment of marker genes across the three sub-clusters of immature granule cells (GC) E) UMAP plots separated by group reveal differential numbers of cluster 3 immature GC in stimulation group. F) Box plot shows increased proportion of Prox1+ immature GC cells in stimulation vs ambient Ts65Dn mice. Center line represents the median; lower and the upper lines represent the 25^th^ and 75^th^ percentiles, respectively; whiskers indicate the smallest and largest values in the 1.5x interquartile range. G-H) Increased cell proliferation in the dentate gyrus of stimulation vs ambient Ts65Dn mice tested via immunolabeling for G) Ki67 (ambient = 6 mice, stimulation = 4 mice); Scale bar = 10 µm. and H) EdU (ambient = 8 mice, stimulation = 8 mice); Scale bar = 50 microns. Error bar indicates mean ± standard error mean. Two tailed, unpaired t- test. *P<0.05, **P<0.01, ***P<0.001.

Previous studies have leveraged snRNA-seq datasets, specifically the transcriptional profile of granule cells (GCs), to address adult neurogenesis in mice [42] and in humans [43]. To investigate whether gamma stimulation can affect adult neurogenesis, we sorted nuclei based on the expression of Prox1, a pan-GC marker [42,44], and selected GC using an *in silico* approach (Materials and Methods, **Fig S10**). We ordered cells along the GC developmental trajectory (**Fig S11**), identified genes whose expression changed as a function of pseudotime (**Fig S11A**), and then analyzed the effect of gamma stimulation on gene expression changes along this GC developmental trajectory (**Table S8**). GO analysis revealed that 40 Hz stimulation induced differential expression of genes related to biological processes of various themes related to adult neurogenesis (e.g., neuron projection development, synapse organization, regulation of RNA splicing, microtubule-based transport, and cell morphogenesis involved in neuron differentiation) (**Fig S11B**) and that some of these processes overlapped with those reported to be downregulated in immature GCs in aging [43]. When analyzing immature granule cells (GCs) in this study, as further confirmed by the expression of immature GC markers (**Fig S10**), we detected three distinct subclusters (**Fig 3C**). Each subcluster was characterized by the expression of specific marker genes that differentiated it from the others (**Fig 3D**). Among these clusters, cluster 3 was transcriptionally distinct from the other two clusters (**Table S9**), with its differentially expressed genes associated with adult neurogenesis-related biological processes (e.g., neuron migration, regulation of neurogenesis, oxidative phosphorylation, and cellular respiration) (**FigS11C)**. Intriguingly, the proportion of cluster 3 was significantly increased in the stimulation group (**Fig 3E-F)** compared to the ambient group, suggesting a potential increase in neurogenesis.

To experimentally validate the impact of gamma stimulation on adult neurogenesis, we adopted two different experimental approaches for directly testing for cell proliferation in the dentate gyrus. In the first approach, fixed brain slices from both the stimulation and ambient groups were stained for Ki67, a marker for cellular proliferation (**Fig 3G**). The stimulation group exhibited a significantly higher number of Ki67+ cells in the sub-granular zone of the dentate gyrus compared to ambient group (stimulation = 18.3+/-1.3, ambient = 12.0+/-4.4, p = 0.027, **Fig 3G**), suggesting increased neurogenesis. In the second approach, a separate cohort of stimulation and ambient Ts65Dn mice were injected with EdU (25 mg/kg), a nucleoside analog of thymidine that labels newly synthesized DNA and thus proliferating cells, for 2 days prior to perfusion (**Fig 3H)**. In this cohort, mice in the stimulation group showed a significant increase in the number of cells positive for EdU in the dentate gyrus (stimulation = 8.1+/-3.1, ambient = 4.5+/-2.9, p = 0.03, **Fig 3H**), again indicating that 40 Hz sensory stimulation increased adult neurogenesis in TsD65n mice.

Together, our results demonstrate that 40Hz stimulation can increase adult neurogenesis and improve cognitive performance in the Ts65Dn mouse model. Furthermore, our findings suggest that these improvements are related to changes in the expression of hippocampal gene regulatory networks that are involved in synaptic organization within the hippocampus and show association with aging and AD and to enhanced adult neurogenesis.

### Discussion

In this study, we investigated the efficacy of a non-invasive multisensory gamma stimulation to improve cognitive functions in male Ts65Dn mice, a widely used mouse model of DS, at 6-8 months of age. The Ts65Dn mice at that age do not exhibit increased levels of APP in the hippocampus [45] and lack the insoluble amyloid plaques and neurofibrillary tangles characteristic of AD [15,46], but do show some early symptoms of AD-related pathologies such as increased levels of soluble amyloid Aß40 and Aß42 in the hippocampus [47], signs of hippocampal-dependent cognitive impairments [46,48], and hippocampal degeneration [49]. Since approximately 90% of adults with DS will develop AD in their lifetimes and show substantial Aß accumulation and NFTs in the brain by age 40 [50], as well as increased soluble Aß in the brain [51], the presence of some AD-like features in Ts65Dn mice makes this model particularly suitable for studying early AD pathology in DS and testing the therapeutic potential of multisensory gamma stimulation. Our study reveals beneficial effects of multisensory gamma stimulation on cognitive performance in hippocampal-dependent tasks and on adult neurogenesis and provides hippocampal gene expression signatures and gene regulatory networks at the single cell resolution. We show that three weeks of daily one-hour multisensory stimulation improved spatial memory and object recognition in Ts65Dn mice. Importantly, these changes were not confounded by locomotory behaviors, suggesting gamma stimulation-specific effects on hippocampal-dependent memory and recognition, which is consistent with previous observations in mouse models of AD and neurodegeneration [6,7].

The specific impact of the sensory gamma stimulation on transcriptomic changes and gene regulatory network within the hippocampus, particularly at the single cell level, has not previously been investigated. Here, we used snRNA-seq of hippocampal tissue from Ts65Dn mice to describe the hippocampal molecular signatures underlying the spatial cognitive benefits from sensory gamma stimulation. We showed that the stimulation increased expression of genes related to synapse organization in excitatory neurons. Moreover, we identified specific gene regulatory modules upregulated by 40 Hz stimulation within the excitatory neurons that significantly overlap with genes downregulated during aging and in AD. This finding further highlights the relevance of these gene modules for cognitive benefits. Sensory-evoked modulation of these genes are particularly intriguing given that early clinical studies have shown sensory 40 Hz stimulation – which is non-invasive, inexpensive, and easily administered in a home setting – to be safe and tolerable in humans [11]. Several of the hub genes in the gene modules with increased expression after the sensory treatment have been implicated in learning and memory, general cognitive function, and/or AD based on GWAS studies, such as *Adcy8*, *Gria4* or *Reln*. A potential role of Reelin in providing cognitive benefits in DS mice is further supported by experimental confirmation. In our previous work, where we identified *Reln*+ neurons as particularly vulnerable in AD [52]. Excitingly, in this study, sensory gamma stimulation increased the number of *Reln+* neurons in the hippocampus in a subregion-specific manner. Given that DS is a significant risk factor for AD in humans, these findings in the Ts65Dn mouse model are particularly intriguing, as the genes with altered expression following 40 Hz stimulation overlap with those implicated in AD, providing potential avenues for treating AD pathologies in DS.

We further identified *TCF4*, a critical factor in adult neurogenesis [40,41], as one of the upstream regulators for the most conserved gene module and not only observed a notable increase in *TCF4* expression specifically in the dentate gyrus after 40 Hz stimulation, but directly demonstrated an increase in adult neurogenesis in the dentate gyrus of Ts65Dn mice. Although the exact role of adult hippocampal neurogenesis on memory continues to be elucidated, extensive evidence has shown its connection to hippocampus dependent tasks both under conditions that deplete neurogenesis [53] as well as conditions that promote neurogenesis (e.g., genetic [54] or pharmacological [55] interventions, exercise [56], and environmental enrichment [57]). In Ts65Dn mice, specifically, interventions that rescue neurogenesis have been shown to lead to improvements in novel object recognition and novel object location [58–61].

Additionally, we noted that 40 Hz stimulation increased expression of genes related to synaptic organization and the number of putative synapses, as identified by co-localization of PSD95 puncta with pre-synaptic synaptophysin puncta, in the dentate gyrus. It is likely that these hippocampal synaptic changes along with increased neurogenesis underlie the improved hippocampal-dependent tasks in 40Hz stimulation.

The present study reveals a beneficial impact of three-week long 40 Hz multisensory gamma stimulation of adult Ts65Dn male mice on their cognitive performance, neurogenesis, and hippocampal synaptic organization highlighting the potential of gamma stimulation as a therapeutic strategy for cognitive deficits in individuals with DS. Of note, one limitation of this study is that the Ts65Dn mice display trisomy for only approximately two-thirds of the genes orthologous to human chromosome 21 (Hsa21), but also for some genes that are not triplicated in human DS, including ∼35 protein-coding genes arising from mouse chromosome 17 (Mmu17) [62,63]. Although the chronic multisensory gamma stimulation did not cause expression changes for these genes, this additional triplication may introduce phenotypic effects unrelated to DS, potentially confounding the interpretation of results specific to DS-related pathology. Further, the transmission of the trisomy is carried through the maternal germline, requiring that the mothers be trisomic, which is generally not the case in humans. Thus, findings in this study should ideally be validated in other alternative DS mouse models [63]. Another limitation is that the study focused solely on hippocampal gene expression changes, even though the prefrontal cortex is a critical brain region involved in spatial memory, particularly for the alternation tests performed in this study. Thus, future research should also investigate the effects on the gene expression changes in the prefrontal cortex. Furthermore, the current study examined the effects of gamma stimulation at 40 Hz exclusively in male mice, highlighting the need for future studies to include both sexes and test other frequencies. Such studies will provide a more comprehensive understanding of the beneficial effects of the multisensory gamma stimulation and to explore potential of its sex-specific effects on cognitive performance [64], hyperactivity [15,65], developmental milestones [66] and soluble amyloid pathology (e.g., Aß40, Aß42) [47] in Ts65Dn mouse model. Additionally, the current study assessed only the short-term memory effects of multisensory gamma stimulation. Future studies should explore its impact on long-term memory, investigate whether multimodal sensory integration is critical for cognitive benefits and assess if treatment benefits can be further increased by longer durations of multisensory 40 Hz treatment and/or earlier treatment onset, such as in the early post-natal stage, or addition of other sensory modalities (e.g., tactile stimulation [67]).

### Materials and Methods

#### Animals

All animal work was approved by the Committee for Animal Care of the Division of Comparative Medicine at the Massachusetts Institute of Technology and by the Institutional Animal Care (MIT- CAC, approval number: 0621-033-24). Male Ts65Dn mice aged 6-8 months were used for all experiments. Mice were housed in groups no larger than five on a standard 12-h light/12-h dark cycle; all experiments were performed during the light cycle. Food and water were provided without restriction. For all experiments, mice from the same litter were divided into different conditions, respectfully. If additional groups were added, respective ambient groups were always repeated concurrently. For tissue collection, mice were anesthetized with isoflurane and cardiac perfused with ice cold DPBS (Thermo Fisher Scientific, 14190235).

#### Concurrent 40 Hz auditory and visual stimulation protocol

Light flicker stimulation was delivered as previously described [10,68]. Mice were transported from the holding room to a separate stimulation room, located on another floor. Mice were habituated under dim light for 30 min before the start of the experiment, and then introduced to the stimulation cage (like the home cage, except without bedding and three of its sides covered with black sheeting). All sensory gamma stimulation protocols were administered daily for 1h/d for the number of days as specified. Mice were exposed to one of two stimulations: dark/quiet or concurrent 40 Hz light flicker and auditory tone for one hour per day. Mice were allowed to freely move inside the cage but did not have access to food or water during the 1-hour stimulation period. An array of light emitting diodes (LEDs) was present on the open side of the cage and was driven to flicker at a frequency of 40 Hz with a square wave current pattern and 50% duty cycle using an Arduino system. The luminescence intensity of light that covered inside the total area of multisensory stimulation cage varied from ∼ 200 – 1000 lux as measured from the back and front of the cage (mice were free to move in the cage). For audio stimulation, a 10 kHz tone at 60 dB was played at 40 Hz with a 4% duty cycle from a speaker located above the stimulation cages. After 1h of stimulation, mice were returned to their home cage and allowed to rest for a further 30 min before being transported back to the holding room. Ambient mice group underwent the same transport and were exposed to similar cages with similar food and water restriction in the same room but experienced only normal room light and natural environmental noise for the 1h duration.

#### Behavior

Behavioral protocols were conducted towards the end of the 3-weeks stimulation protocol. Ethanol was used to wipe down all testing apparatuses between uses.

##### Novel Object Recognition

The novel object recognition (NOR) task consisted of a habituation phase followed by training and testing performed the following day, as previously described [69]. 24 hours before training, mice were habituated to an open testing arena (40 cm L x 40 cm W x 35 cm H) for 10 min, during which total distance (cm), time in the center (s), and velocity (cm/s) were calculated (TSE Systems) to evaluate open field behavior. During training, mice were placed into the same box with two identical objects placed in opposite corners. Mice were allowed a total of 20 s of object interaction time (within a maximum time frame of 10 minutes), and then immediately removed from the arena. Object memory was tested 1 hour later using the same procedure during training, except one object was replaced with a novel one in its place. Object exploration was recorded when the snout contacted either object and was calculated by a recognition index, RI = (Tnovel/(Tnovel+Tfamiliar)) x100, where Tnovel and Tfamiliar indicate the time spent with the novel and familiar object, respectively.

##### Novel Object Location

The novel location recognition (NOL) task was performed using the same procedure as the object recognition task, except two identical objects were used for both training and testing, and one object was displaced to a novel location during testing.

#### Spontaneous Alternation

A Y-maze was used for testing with an apparatus with three equal arms (30 cm length, 10 cm width, and 20 cm height) placed 120° apart, made of opaque acrylic glass. A mouse was placed at the maze center and was allowed 7-min of exploration of the environment. An arm entry was scored when the mouse entered the arm with all four paws. Total number of entries (N) and number of ‘correct’ triplets (M, consecutive choices of each of the three arms without re-entries) was evaluated. The alternation rate was computed according to the formula: R (%) = (M/(N-2))x100.

#### Single-nucleus RNA sequencing

24 hours after the final stimulation session, the hippocampi of the mice were dissected and snap frozen in liquid nitrogen and stored at −80 °C. The protocol for the isolation of nuclei from frozen brain tissue was adapted from a previous study [70]. All procedures were carried out on ice. Briefly, two mouse hippocampi were pooled per sample (3 samples per condition) and homogenized in 1 ml homogenization buffer (320 mM sucrose, 5 mM CaCl2, 3 mM Mg (CH3COO)2, 10 mM Tris HCl pH 7.8, 0.1 mM EDTA pH 8.0, 0.1% IGEPAL CA-630, 1 mM β-mercaptoethanol, and 0.4 U µl-1 recombinant RNase inhibitor (Clontech) using a Wheaton Dounce tissue grinder. The homogenized tissue was filtered through a 40-μm cell strainer, mixed with an equal volume of working solution (50% OptiPrep density gradient medium (Sigma-Aldrich), 5 mM CaCl2, 3 mM Mg(CH3COO)2, 10 mM Tris HCl pH 7.8, 0.1 mM EDTA pH 8.0, and 1 mM β-mercaptoethanol) and loaded on top of an OptiPrep density gradient (29% OptiPrep solution (29% OptiPrep density gradient medium,134 mM sucrose, 5 mM CaCl2, 3 mM Mg(CH3COO)2, 10 mM Tris HCl pH 7.8, 0.1 mM EDTA pH 8.0, 1 mM β-mercaptoethanol, 0.04% IGEPAL CA- 630, and 0.17 U µl-1 recombinant RNase inhibitor) on top of 35% OptiPrep solution (35% OptiPrep density gradient medium, 96 mM sucrose, 5 mM CaCl2, 3 mM Mg(CH3COO)2, 10 mM Tris HCl pH 7.8, 0.1 mM EDTA pH 8.0, 1 mM β-mercaptoethanol, 0.03% IGEPAL CA-630, and 0.12 U µl-1 recombinant RNase inhibitor). The nuclei were separated by ultracentrifugation using an SW32 rotor (5 min, 10,000xG, 4°C). Nuclei were collected from the 29%/35% interphase, washed with PBS containing 0.04% BSA, centrifuged at 300g for 3 min (4 °C), and washed with 1 ml of PBS containing 1% BSA. The nuclei were counted and diluted to a concentration of 1,000 nuclei per microliter in PBS containing 1% BSA. Libraries were prepared using the Chromium Single Cell 3′ Reagent Kits v.3.1 (Dual Index) according to the manufacturer’s protocol (10X Genomics, Pleasanton, CA). The snRNA-seq libraries were sequenced using NextSeq 500/550 High Output (150 cycles).

#### Single nuclei RNA-seq data analysis

##### Data preprocessing, differential expression, gene ontology analyses

Gene counts were retrieved by mapping raw reads to the mouse genome (mm10) using CellRanger software (v 7.0.1). Initial preprocessing of the samples (e.g., removal of ambient noise, identification and removal of contamination from adjacent brain regions, and stressed or damaged cell and low quality cells) were performed using ACTIONet [71]. Downstream analysis was performed in Seurat (v 4.3.0). Nuclei with a minimum 500 unique molecular identifiers (UMIs) and less than 5% mitochondrial genes were considered for downstream analysis. Doublets were identified and removed using scds (v 1.10.0) and genes with no expression was discarded. Count data were log-normalized using NormalizeData and top 2000 highly variable genes were determined using FindVariableFeatures. Expression of these genes were subsequently used to scale gene expression to zero mean and unit variance with ScaleData function. Scaled data was further regressed out for mitochondrial percentage. Top 75 principal components were estimated using RunPCA and aligned with Harmony to account for technical batch effects considering individual sample as batch. Optimum number of principle components (PCs) were determined based on the ElbowPlot and the first 16 PCs was used as input for downstream analyses. A neighborhood graph was constructed based on the Euclidean distance metric in the adjusted PC space using FindNeighbors, and nuclei were clustered using the Louvain method in FindClusters at resolution 0.2. RunUMAP functions with min.dist = 0.5 and spread = 0.5 were used to calculate two-dimensional UMAP coordinates and nuclei clusters were visualized using DimPlot. Cell type specific clusters were annotated using canonical markers. Nuclei clusters expressing mixed cell type specific markers were discarded. Cell type specific marker genes were identified based on the differential expression of each cell type against all other cell types using FindMarkers and genes with significance (adjusted P value<0.05) and increased expression were only considered for cell type specific markers. Differential gene expression analysis was performed using MAST (v 1.20.0). Genes with log2 fold change >0.1 and adjusted p value <0.05 were defined as differentially expressed. Module score was calculated using AddModuleScore in Seurat. Gene ontology analyses were performed using either Gene ontology (http://geneontology.org) or Metascape (https://www.metascape.org/). For supervised GO module analysis particularly for oxidative phosphorylation (GO:0006119) and cellular respiration (GO:0045333), we retrieved genes related to these GO terms using gconvert function of gprofiler2 (v 0.2.1). Co-expression GO module score was calculated using AddModuleScore function in Seurat.

##### Weighted gene co-expression analysis

Weighted gene co-expression analysis (WGCNA) was performed as previously described in [31,72,73]. Briefly, “metacells” were constructed computing mean expression from 25 neighbors using k-nearest neighbors. Normalized and scaled metacells were then used to calculate pair-wise correlations between all gene pairs. Subsequently, based on the approximate scale-free topology, a threshold power of 4 was selected to emphasize the stronger correlations and to compute pair-wise topological overlap to construct a signed network. Modules with minimum size of 25 and deep split of 4 were used for the analysis. Closely related modules were merged using dissimilarity correlation threshold of 0.2. Modules were summarized as network of modular eigengenes (MEs), which was defined based on the first major component of the module and used to compare between the two groups. Module membership of genes was defined as the correlation of gene expression profile with MEs. Hub genes were defined based on the intra-modular connectivity (kME) and top 25 hub genes were plotted using igraph package (v 1.3.1). Association between hub module genes and transcription factors was analyzed using ChEA3 [74].

##### Analyses of granule cells

To select granule cells (GCs) we first filtered the snRNA-seq data based on the expression level of Prox1, which is a pan GC marker. Only cells with an expression level of Prox1 greater than 1 were retained for subsequent analyses. Cells were normalized, scaled, harmonized and top 30 components were used to construct cell clusters and UMAP at resolution of 0.8. Cluster specific marker genes were defined using FindAllMarkers (only.pos = TRUE, min.pct = 0.25, assay = “RNA”) and top 200 genes were used to perform gene enrichment analyses with granule cell specific markers defined in the previous study [75]. Based on the enrichment results, cell clusters were annotated into glial (radial glial), and granule cells (neuroblasts, mature and immature). Clusters that did not show overlap to the marker genes were labeled as unknown cluster type. Annotated granule cells (immature, mature, neuroblasts) were considered for downstream analyses and pseudotime analysis was performed using Monocle. Briefly, we kept the original clustering and dimension reduction from Seurat and created a Monocle cell data set (CDS) using the new_cell_data_set function and treated all cells as a single partition. For trajectory analysis, learn_graph function was used to learn the trajectory graph of the cells. Cells were subsequently assigned a pseudotime value based on their projection on the principal graph learned in the learn_graph function and order_cells function was used to order cells along the trajectory. To find genes whose expression changes along the pseudotime, we then applied a generalized additive model along the principal trajectory via graph_test function. Genes with q-value less than 0.05 were considered only for downstream analysis. Top 100 pseudotime associated genes were visualized as heatmap using Heatmap function. Genes that are differentially expressed between ambient and stimulation groups along the trajectory were determined using fit_models with q- value cutoff set at 0.05. Gene ontology analysis was performed using Metascape and ClusterProfiler.

##### Analyses of published datasets

Hippocampal bulk RNA-seq raw data pertaining to 4, 8, 12 and 18 months of wild type mice were retrieved from GSE168137 and analyzed. Single cell RNA-seq data from frontal cortex and striatum of 4 weeks and 90 weeks of mice was retrieved from GSE207848 and processed in house. List of downregulated genes in excitatory neurons of old individuals compared to young subjects were retrieved from [76]. Cortical and hippocampal bulk RNA-seq data related to Ts65Dn and littermate controls were downloaded from GSE213500 and processed in house. List of downregulated genes in the excitatory neurons of prefrontal cortex from patients with Down syndrome was retrieved from the provided supplementary file in [77]. Bulk RNA-seq data related to 5xFAD and wild type mice of 4 and 8 months were downloaded from GSE168137 and analyzed in house. Single cell data related to 5xFAD and wild type mice of 7 months old were retrieved from GSE140511 and processed in house. Hippocampal excitatory neuronal transcriptomic data from individuals with Alzheimeŕs disease and controls were provided by [52]. For Bulk RNA-seq data, genes with low no expression were removed. Differential expression analysis was performed using DESeq2 (v 1.34.0). Genes with FDR<0.05 were defined as differentially expressed. For single cell data related to aging and AD, data were processed as described above using Seurat and cell clusters were visualized in UMAP space. Cell clusters enriched for Slc17a7/VGLUT1, marker for excitatory neuron, were further analyzed for differential expression analysis. Differentially expressed genes (FDR <0.05, abs(log2FC)>0.1) in the SLC7A7+ human hippocampal snRNA-seq data were determined after adjusting for age, sex, and post-mortem interval. Differential expression analyses were performed using MAST. Module score in bulk RNA-seq data was computed using svd as implemented in moduleEigengene of WGCNA package. Hypergeometric overlap analysis was performed using GeneOverlap package in R.

#### Immunohistochemistry

24 hours after the final stimulation session, mice were transcardially perfused with 40 ml of ice-cold phosphate buffered saline (PBS) followed by 40 ml of 4% paraformaldehyde (PFA) in PBS. Brains were removed and post-fixed in 4% PFA overnight at 4°C and transferred to PBS prior to sectioning. Brains were sectioned 40 μm thick with a Leica VT1000S vibratome (Leica). Sections were permeabilized and blocked in PBS with 0.3% Triton X-100 and 10% donkey serum at room temperature for 2-hrs. Sections were incubated overnight at 4°C in primary antibody containing PBS with 0.3% Triton X-100 and 10% donkey serum. Primary antibodies were anti-synaptophysin (Synaptic Systems, 101004), anti-PSD95 (Abcam, ab18258), anti-Ki67 (Abcam, ab15580), anti-cFos (Santa Cruz Biotech, sc-166940), anti-Reelin (Millipore Sigma, MAB5364), anti-TCF4 (Protein Tech, 13838-1-AP), anti-NeuN (Synaptic Systems, 266004). The following day, brain sections were incubated with fluorescently conjugated secondary antibodies (Jackson ImmunoResearch) for 2 hours at room temperature, and nuclei were stained with DAPI (Invitrogen, D1306). Images were acquired using a confocal microscope (LSM 710; Zeiss or LSM 900; Zeiss).

##### EdU Staining

EdU staining was conducted using the Click-iT™ EdU imaging kit (ThermoFisher, C10340) according to the manufacturer’s protocol. This protocol is normally intended for use in cell culture but was adapted for histological staining of brain tissue. Each mouse was intraperitoneally injected with EdU 25 mg/kg for 2 days prior to perfusion. Slides containing mounted frozen brain sections were allowed to thaw to room temperature and then fixed with 4% paraformaldehyde in phosphate buffer saline (PBS) for 15 min. After washing twice with 3% bovine serum albumin (BSA) in PBS, the sections were permeablized with 0.5% Triton X-100 in PBS for 20 min. The sections were again washed twice with 3% BSA in PBS and then incubated with a Click-iT™ reaction cocktail containing Click-iT™ reaction buffer, CuSO_4_, Alexa Fluor® 594 Azide, and reaction buffer additive for 30 min while protected from light. The sections were washed once more with 3% BSA in PBS before being mounted and imaged.

#### Image analysis

##### c-Fos quantification

Images acquired at 20x magnification and whole hippocampal cFos+ nuclei were analyzed in Imaris (v 10.2). Briefly, surfaces were generated for both cFos and DAPI and cFos+ nuclei were quantified by identifying DAPI+ nuclei overlapping with cFos signals. To account for technical bias, the percentage of cFos+ nuclei was calculated relative to the total number of analyzed nuclei.

##### Quantification of colocalization of synaptophysin and PSD95

Following the image acquisition using 63x objective, files were imported into Imaris (v9.9). First, the Spots feature of the software was utilized to detect PSD95 and synaptophysin puncta. For each fluorescence channel, minimum and maximum intensity values were collected from every image to maximize detection of puncta and minimize background noise. Means calculated from these minimum and maximum intensity values became the intensity thresholds for the Spots applied to every image. The Spots using the cohort’s shared threshold values were applied to one sample image, then saved as a Macro and applied to all other images. After applying Spots, the colocalization function was executed to mark and quantify points from each Spots channel that were within 2um of each other. To validate this proxy method for quantifying mature functional synapses, another method for colocalization analysis was also executed. To do this, the Imaris software’s Colocalization Channel function was used. Minimum and maximum thresholds were once again collected from each fluorescence channel to build the new colocalization channel. After calculating the means of the threshold values in each, the channel was built and manually added to every image file. Spots were then added to the new channel to register and quantify the colocalized points detected. Data was analyzed from both methods of analysis to confirm trend. To quantify Reln+ neurons, brain slices were immunostained with anti-Reelin, anti-NeuN and DAPI and images were acquired using the 20x objective in LSM 900. Image analysis was performed using Imaris (v9.9 and v10.2). First, we created surfaces for Reln and NeuN individually. Then, we counted the number of Reln surfaces that overlapped with those of NeuN to identify Reln+ neurons. We calculated the ratio of Reln+ neurons to the total number of neurons to determine the proportion of Reln+ neurons.

To investigate TCF4 expression changes between the two groups, we performed immunostaining experiment using anti-TCF4 and DAPI on brain slices and acquired images (20x, LSM900) from the dentate gyrus, CA1 and CA3 of the hippocampus. For TCF4 mean intensity analysis, we used the Measure function in ImageJ software (v2.14.0/1.54f). To minimize the technical bias, we averaged the TCF4 intensity across all replicates for each mouse and normalized it to the mean DAPI signal for all control samples.

#### Statistical Analyses

Statistical analysis was conducted in R (v 4.1.2) or Prism (v 8). Statistical details of the experiments, including the statistical tests used, exact value of n, definition of measures can be found in the corresponding figure legends. Unless otherwise stated, statistical significance was set at 0.05.

## Acknowledgements

We are grateful to the following individuals and organizations for their kind contributions and support of this work: the Alana Down syndrome Center at MIT and the Alana USA Foundation, Inc, National Science Foundation Graduate Research Fellowship under Grant No.1122374, “la Caixa” Banking Foundation and fellowships for DRA, MRI is a EMBO long term postdoctoral fellow, MM was supported with fellowships from Barbara J. Weedon, Henry E. Singleton, the Hubolow family. ESB acknowledges HHMI and Charles Hieken. We are also thankful to Dr Nancy Kopell, Dr Michelle McCarthy and Tsai lab members for their helpful comments during preparation of this manuscript.

## Supplementary Figures

**Fig S1.**
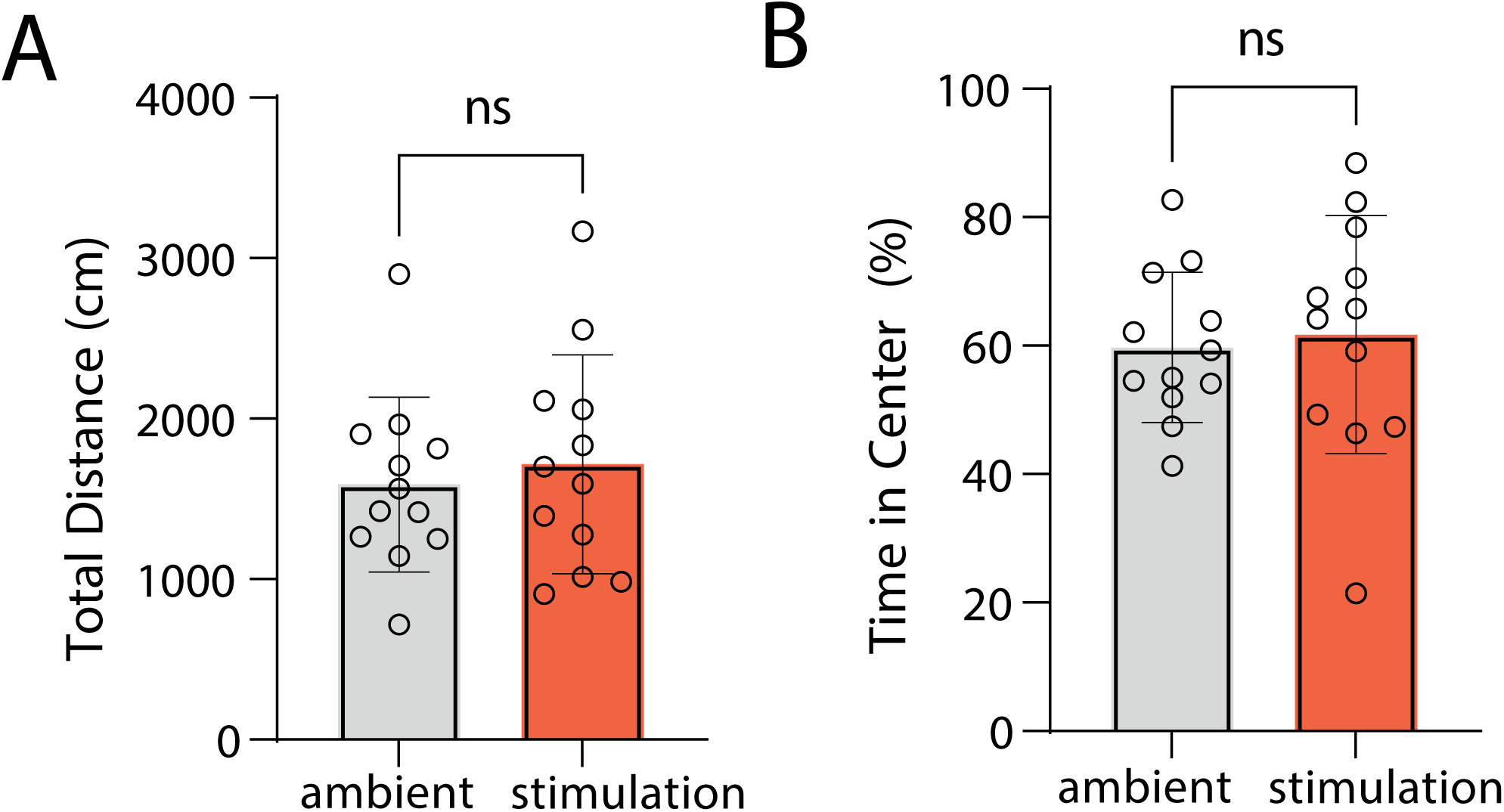
Multisensory gamma stimulation does not affect locomotor and anxiety-like behavior in Ts65Dn mice. A) Quantification of total distance traveled during open field test B) Quantification of time spent in the center. Error bars indicate mean ± standard deviation, unpaired t-test, two-tailed, ambient = 12, stimulation = 12.

**Fig S2.**
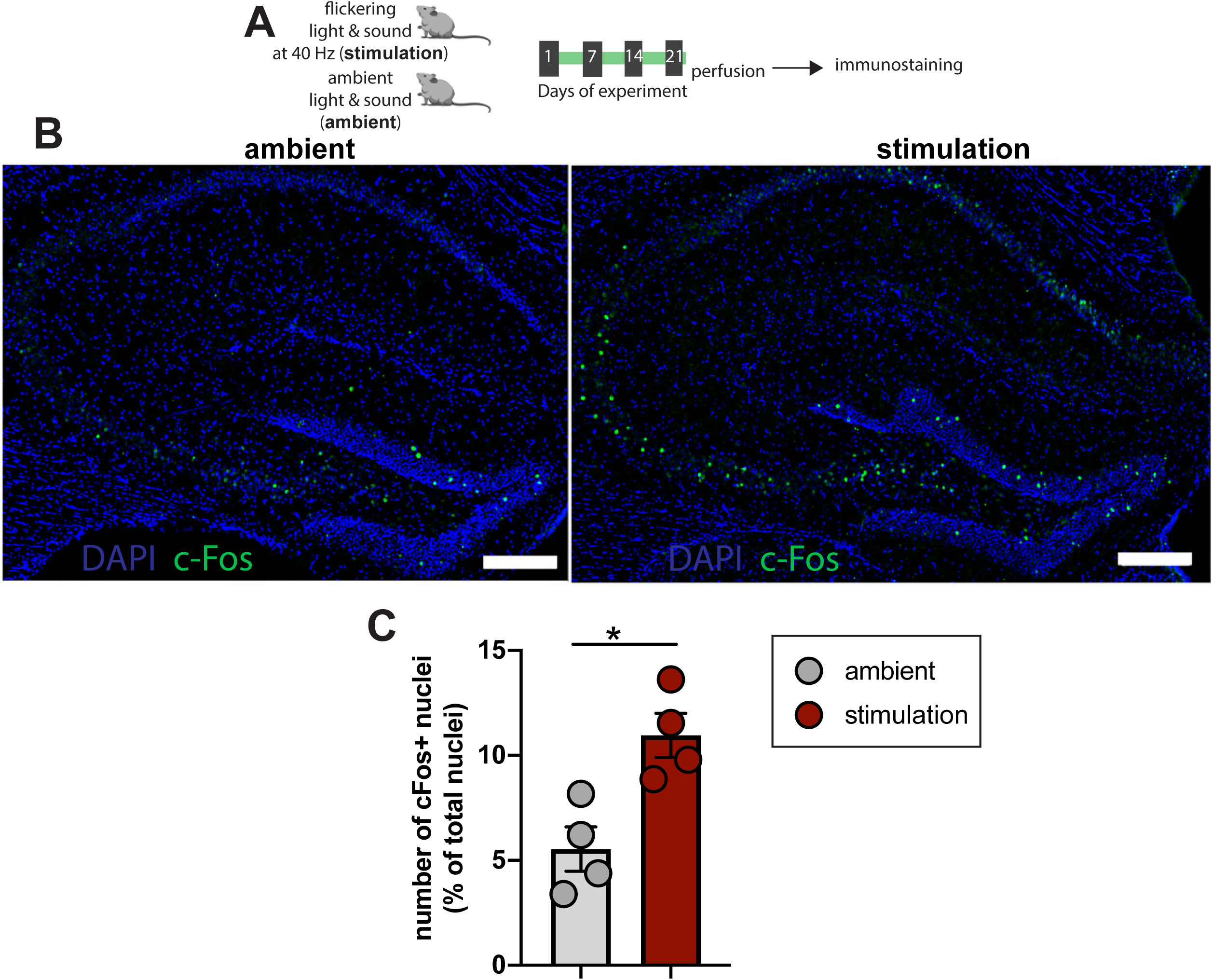
Multisensory gamma stimulation increases c-Fos positive nuclei in the hippocampus of Ts65Dn mice. A) Experimental outline. B) Representative images of c-Fos+ nuclei in the hippocampus after ambient or 40 Hz stimulation. Blue represents DAPI and green represents c-Fos signals. Scale bar = 200 µm. C) Bar plots showing the percentage of c-Fos+ nuclei in the two groups. N = 4 mice /group. Error bar indicates mean ± sem. Two-tailed, unpaired t-test, *P<0.05.

**Fig S3.**
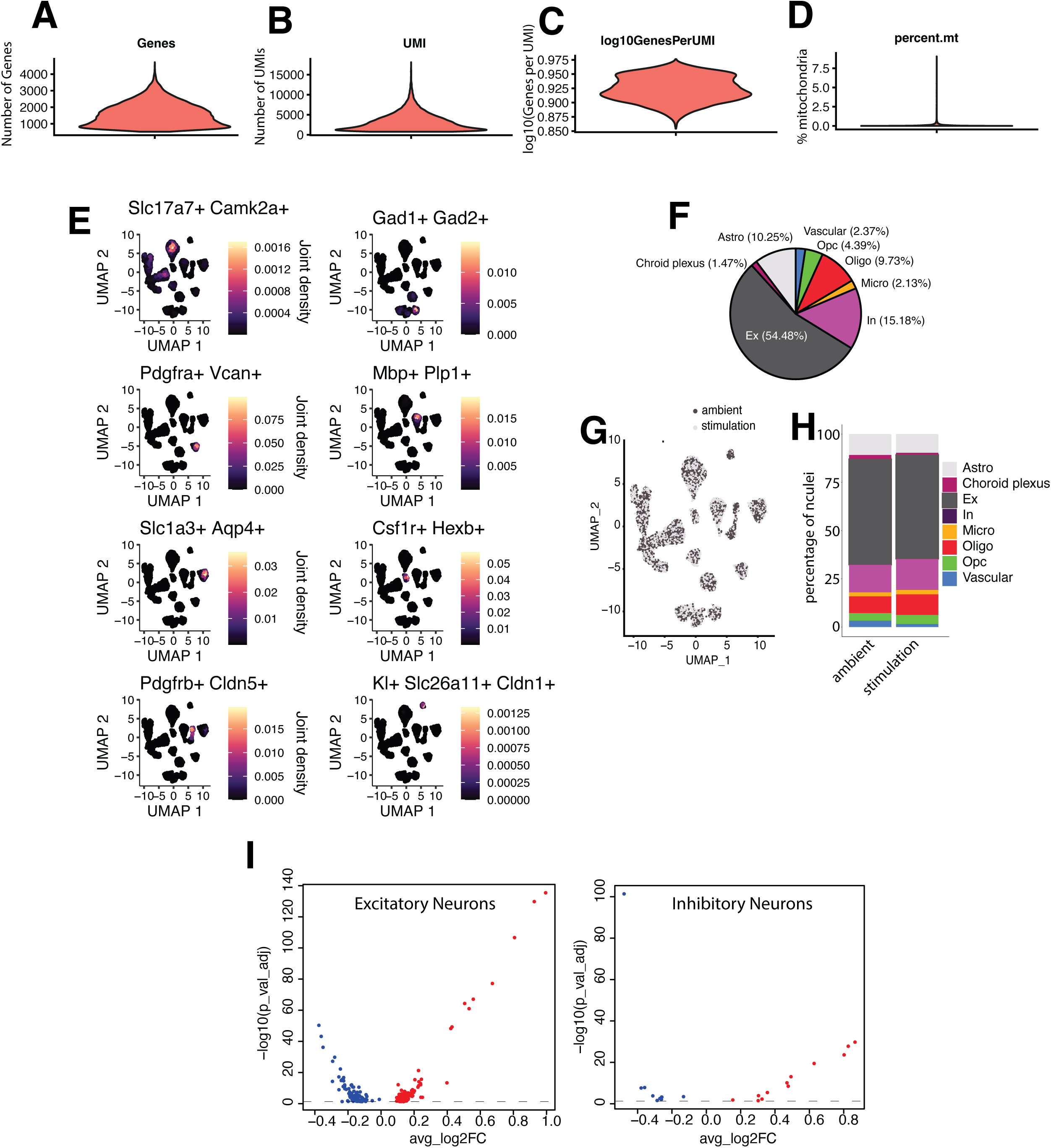
Quality control of the hippocampal snRNA-seq data from Ts65Dn mice. Distribution of A) number of genes B) number of UMI, C) complexity and D) mitochondrial percentage across 15884 nuclei (ambient: 7665, stimulation: 8219). E) Canonical cell type specific marker expression in cognate cell types. Slc17a7, Camk2a in excitatory neurons; Gad1, Gad2 in inhibitory neurons; Pdgfra, Vcan in OPC; Mbp, Plp1 in Oligodendrocytes; Slc1a3, Aqp4 in Astrocytes; Csf1r, Hexb in microglia; Pdgfrb, Cldn5 in vascular cells, KI, Slc26a11, Cldn1 in choroid plexus cells. F) Pie chart displaying the distribution of detected cell types. G) Nuclei from stimulation and ambient Ts65Dn mice groups are well integrated. H) Proportion of nuclei detected across cell types in both groups. I) Volcano plots showing genes that are significantly altered in excitatory and inhibitory neurons in the stimulation group compared to the ambient group.

**Fig S4.**
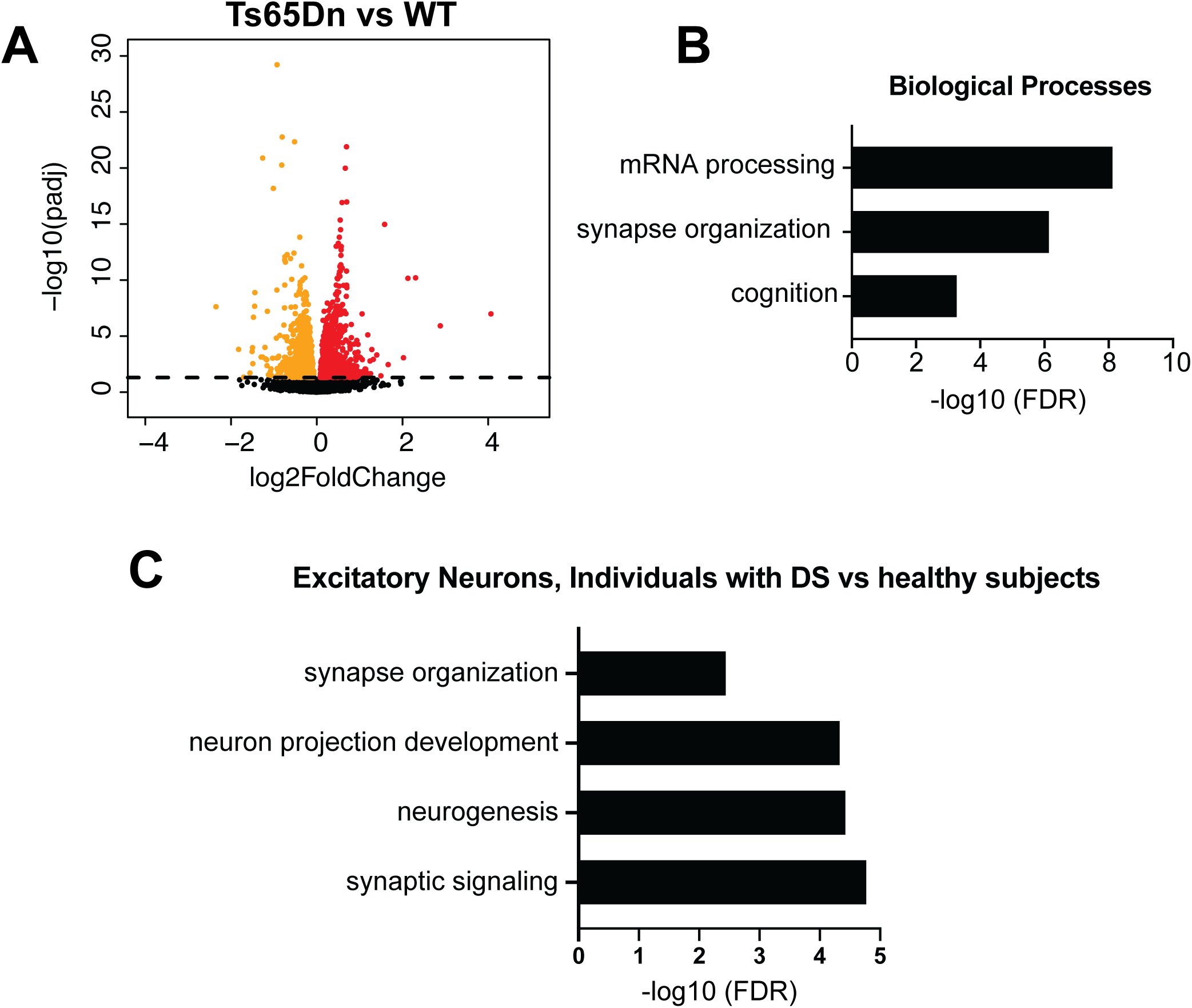
Top biological processes downregulated in Ts65Dn mice and individuals with DS. A) Differential gene expression observed in bulk RNA-seq data from hippocampi and cortices of Ts65Dn mice (n = 6) and littermate controls (n = 6). Data were retrieved from GSE213500. Gene counts were normalized and differential expression analysis was performed using DESeq2 after adjusting for region specific difference. B) Gene ontology analyses revealed that the genes related to mRNA processing, synapse organization and cognition are downregulated in Ts65Dn mice compared to the controls. C) GO analysis of genes downregulated in excitatory neurons of *post mortem* brains of individuals with DS compared to healthy subjects. Gene expression changes data were retrieved from Palmer CR et al^61^.

**Fig S5.**
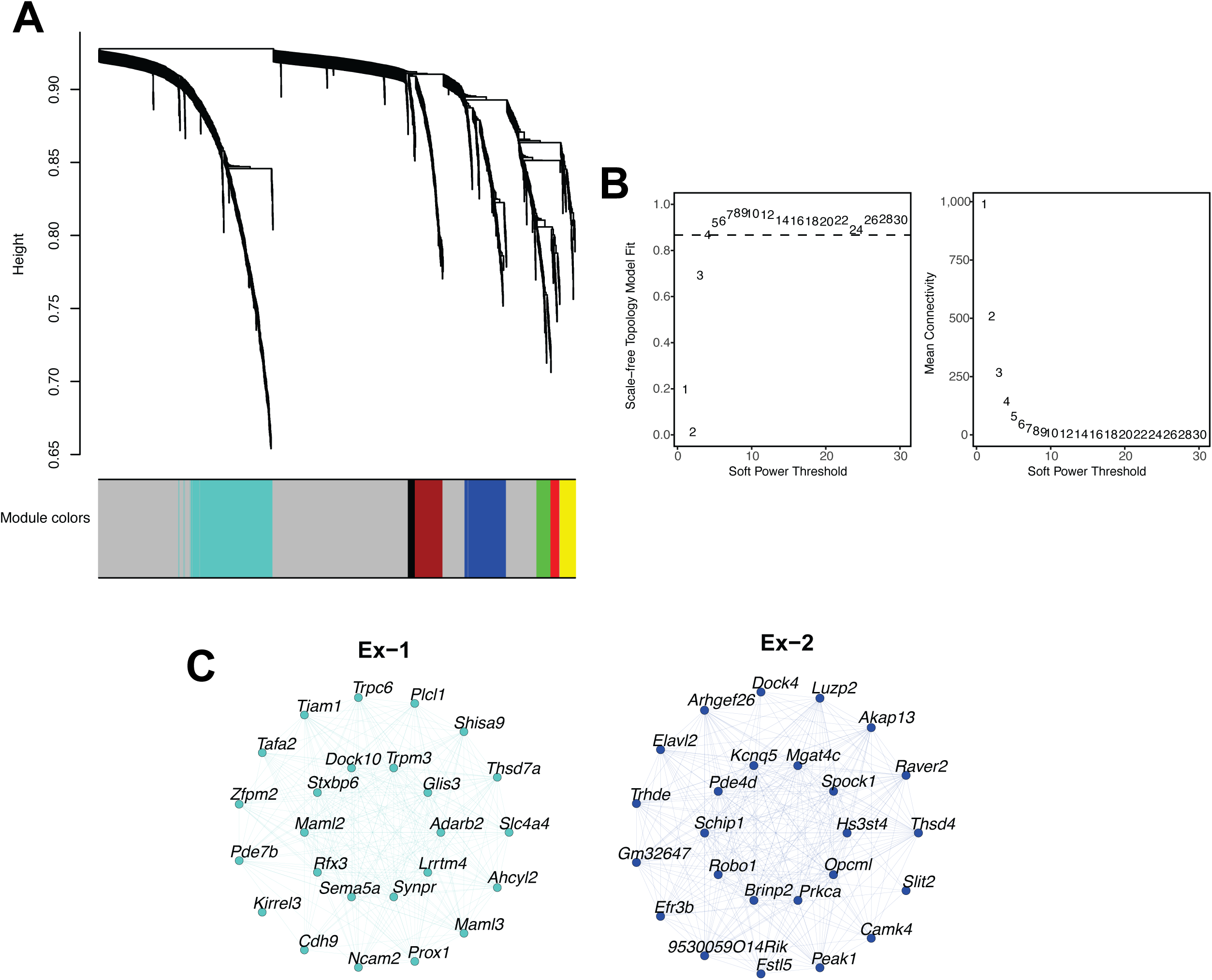
Weighted gene co-expression analysis. A) Cluster dendrogram for seven excitatory gene modules. B) soft-power threshold was determined based on scale-free topology measure and mean connectivity (see Methods). C) Hub genes of Ex-1 and Ex-2 module.

**Fig S6.**
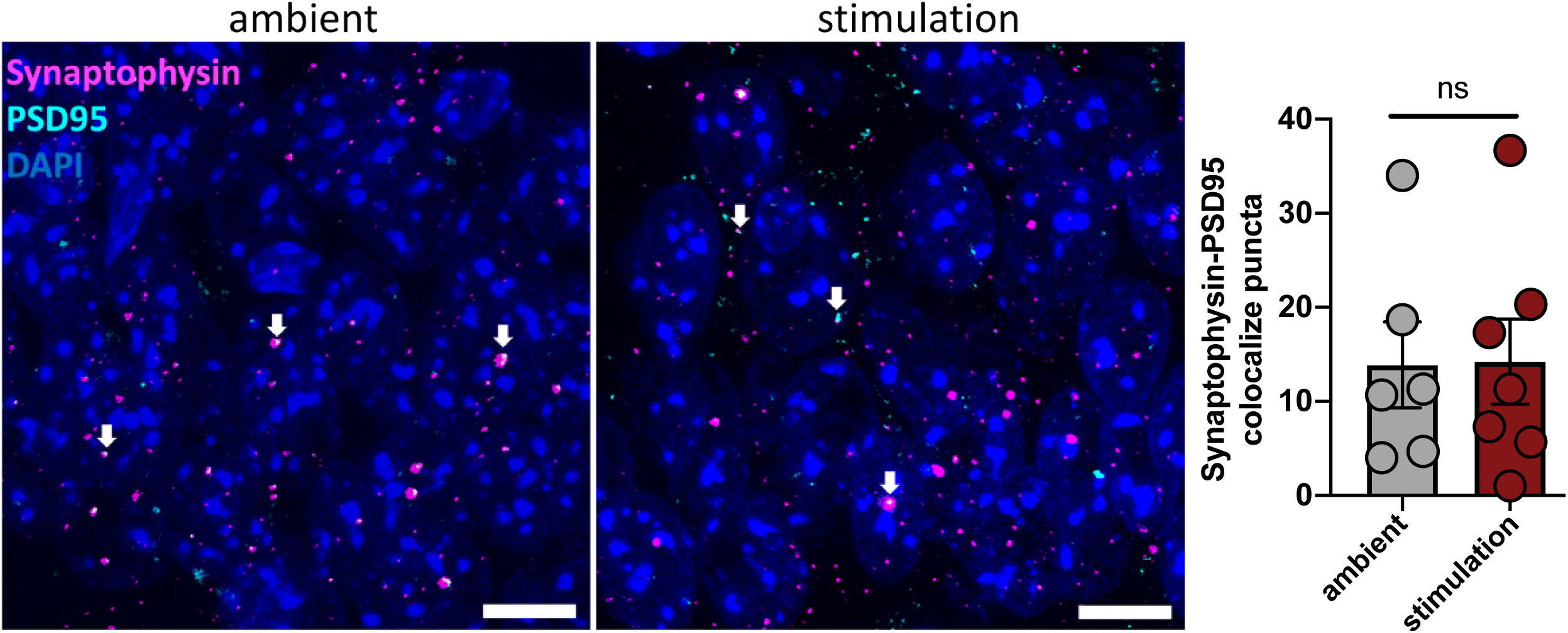
Immunostaining-based analysis of putative synapses in the hippocampal CA1 region between ambient and stimulation groups. (Left) Representative images showing PSD95 and Synaptophysin co-localization. White arrows indicate colocalized puncta. Scale bar = 10 µm. (Right) Bar plots showing the absolute number of colocalized puncta. Statistical significance was determined using a two-tailed, unpaired *t*-test, *N* = 6–7 per group. Error bar: mean ± standard error mean.

**Fig S7.**
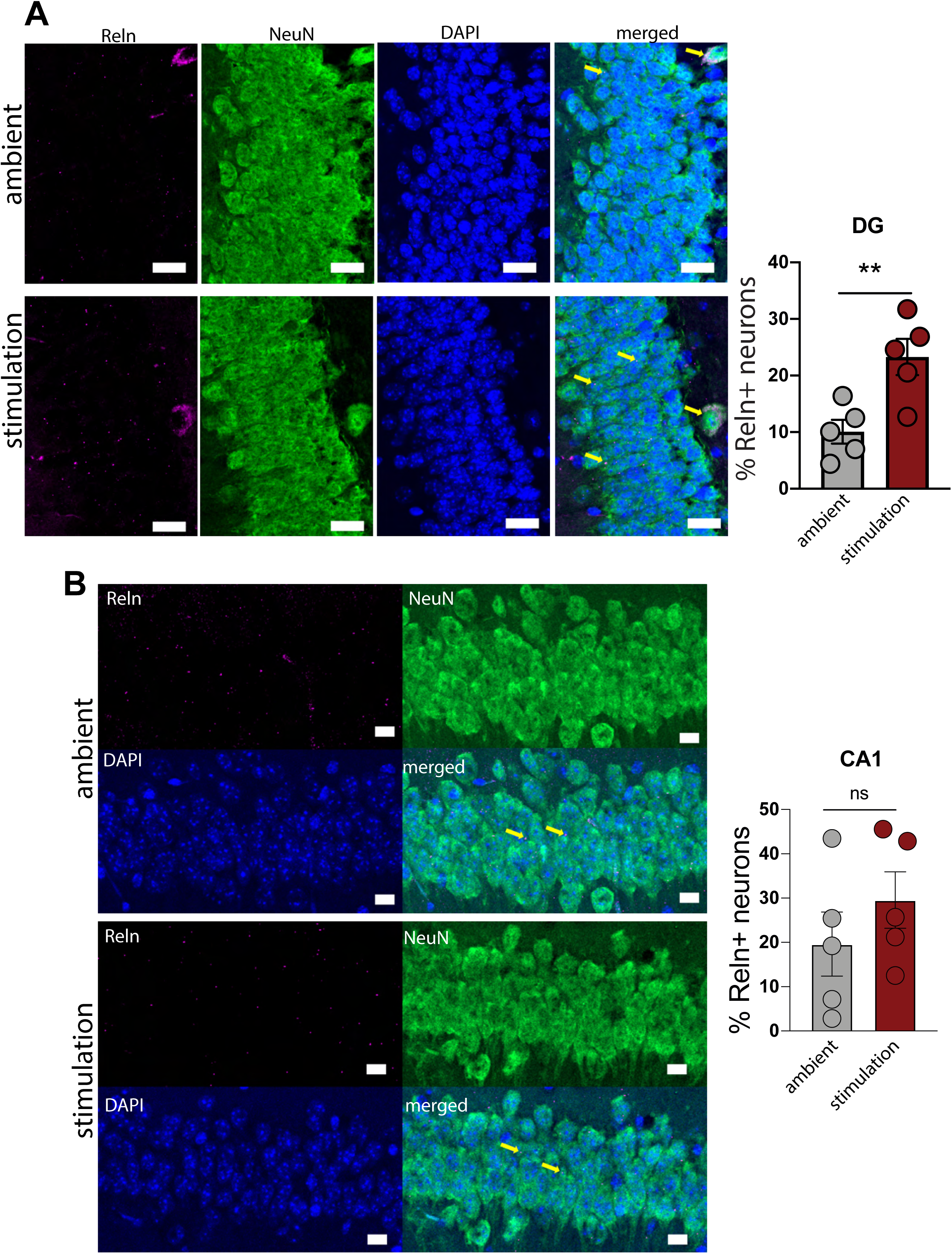
Immunostaining based analyses of Reln+ neurons in hippocampal DG and CA1 regions between ambient and stimulation groups. A) (Left) Representative images of Reln+ neurons in the DG granule cell layer of the hippocampus. (Right) Barplots show percentages of Reln+ neuorns between two groups. 40 Hz stimulation increased percentages of Reln+ neurons in DG of DS mice. Scale bar = 20 µm. Two-tailed t-test, **P<0.01, N = 5 per group. B) (Left) Representative images of Reln+ neurons in the hippocampal CA1 region. (Right) Barplots show percentages of Reln+ neuorns between two groups. Scale bar = 10 µm. Two-tailed, unpaired t-test, N = 5 per group.

**Fig S8.**
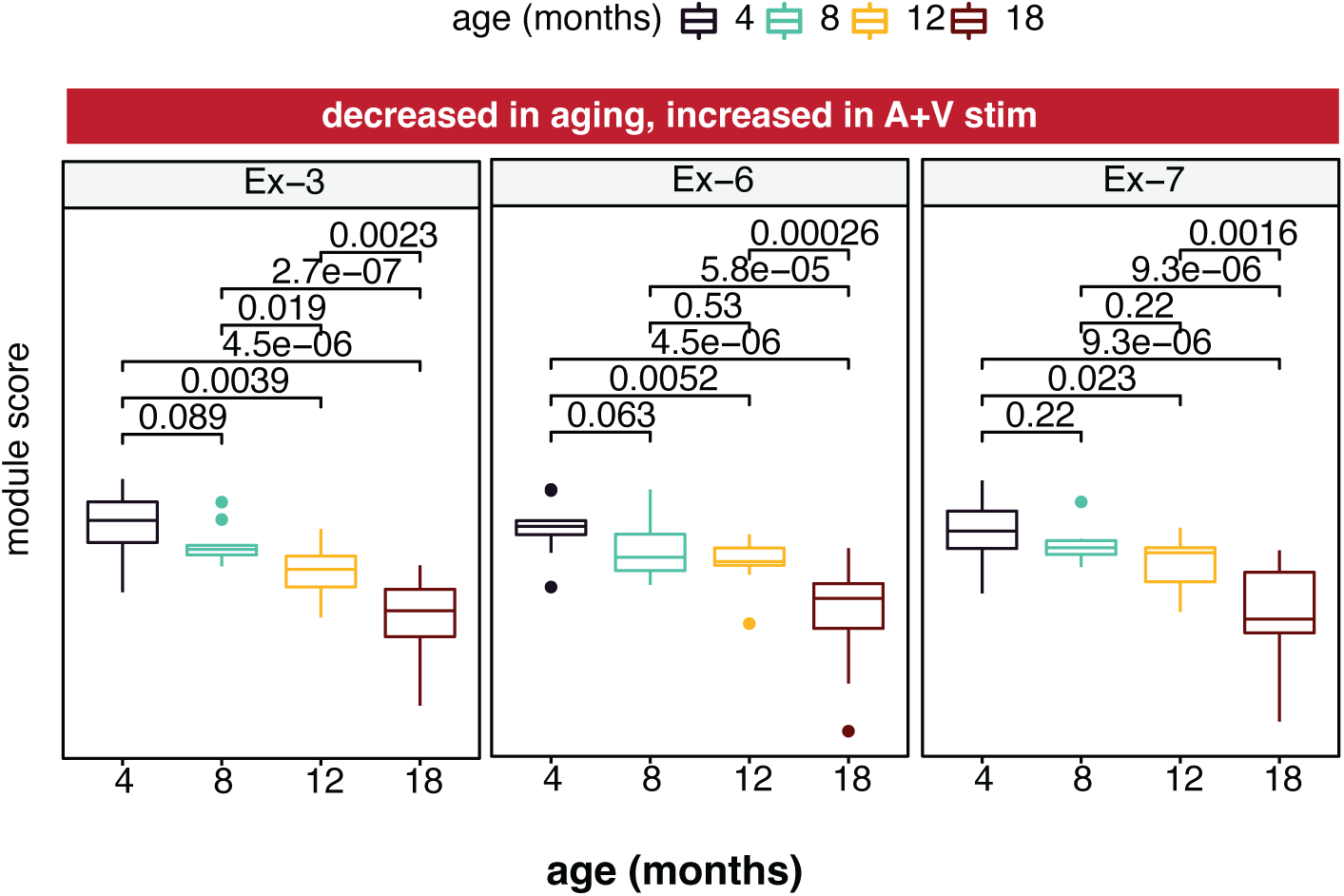
Co-expression pattern of excitatory gene modules along aging. Expression patterns of module Ex-3, Ex-6 and Ex-7 display reduced co-expression at old age (18 months). Boxplots show changes in gene module expression along aging in the hippocampus. Y-axis shows gene module expression eigenvalue. In the box plots, median is marked with the center line, while the lower and the upper lines represent the 25th and 75th percentiles, respectively. The whiskers indicate the smallest and largest values respectively in the 1.5x interquartile range. P-value determined by Kruskal-Wallis test.

**Fig S9.**
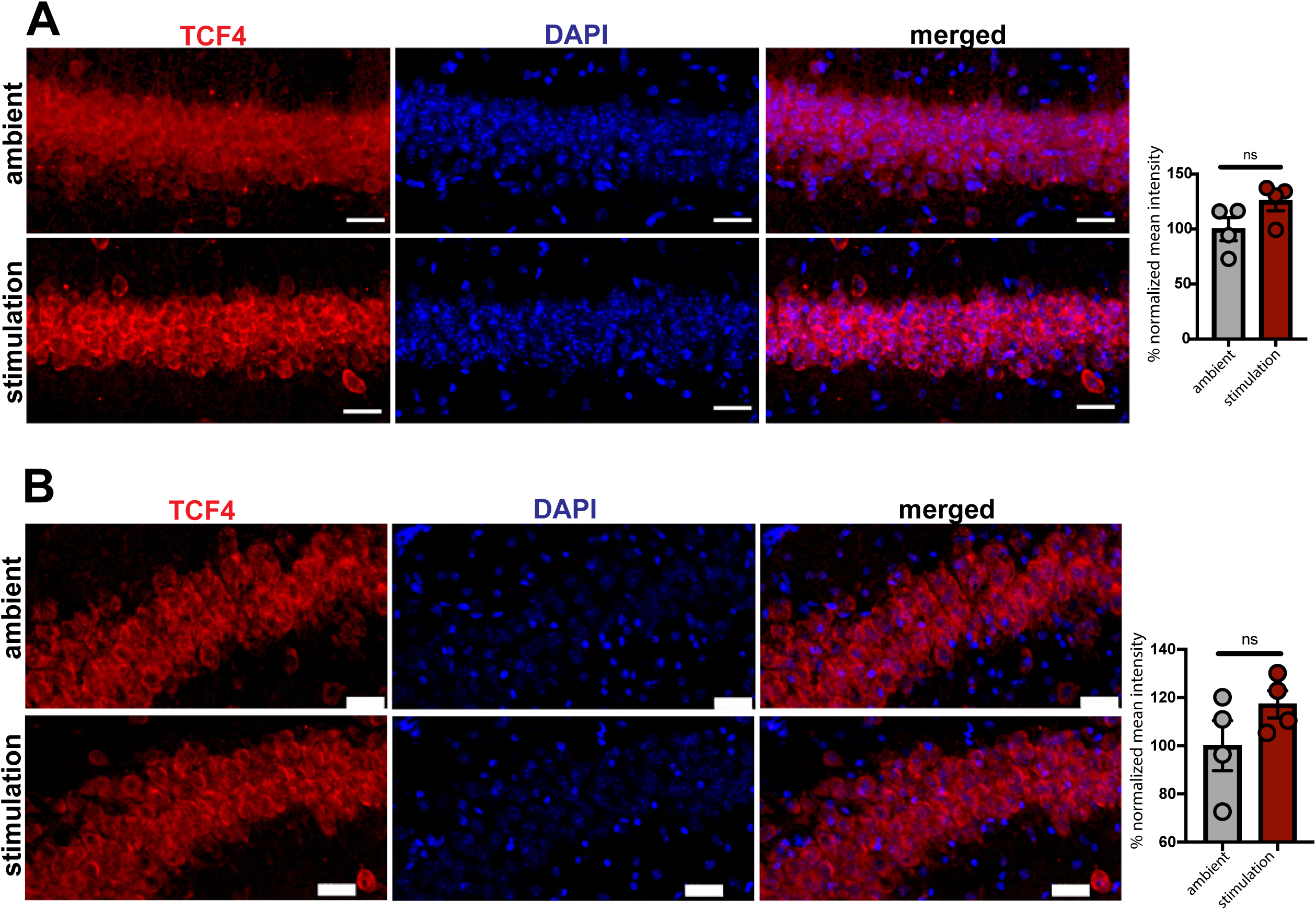
TCF4 expression in the hippocampal CA1 and CA3 between ambient and stimulation groups. A) (Left) Representative images of TCF4 expression in the CA1 region of the hippocampus. (Right) Bar plots showing normalized mean intensity (% of the ambient group) between the two groups. Two-tailed, unpaired *t*-test, N = 4 per group. Scale bar = 30 µm. B) (Left) Representative images of TCF4 expression in the CA3 region of the hippocampus. (Right) Bar plots showing normalized mean intensity (% of the ambient group) between the two groups. Two-tailed *t*-test, N = 4 per group. Scale bar = 30 µm.

**Fig S10:**
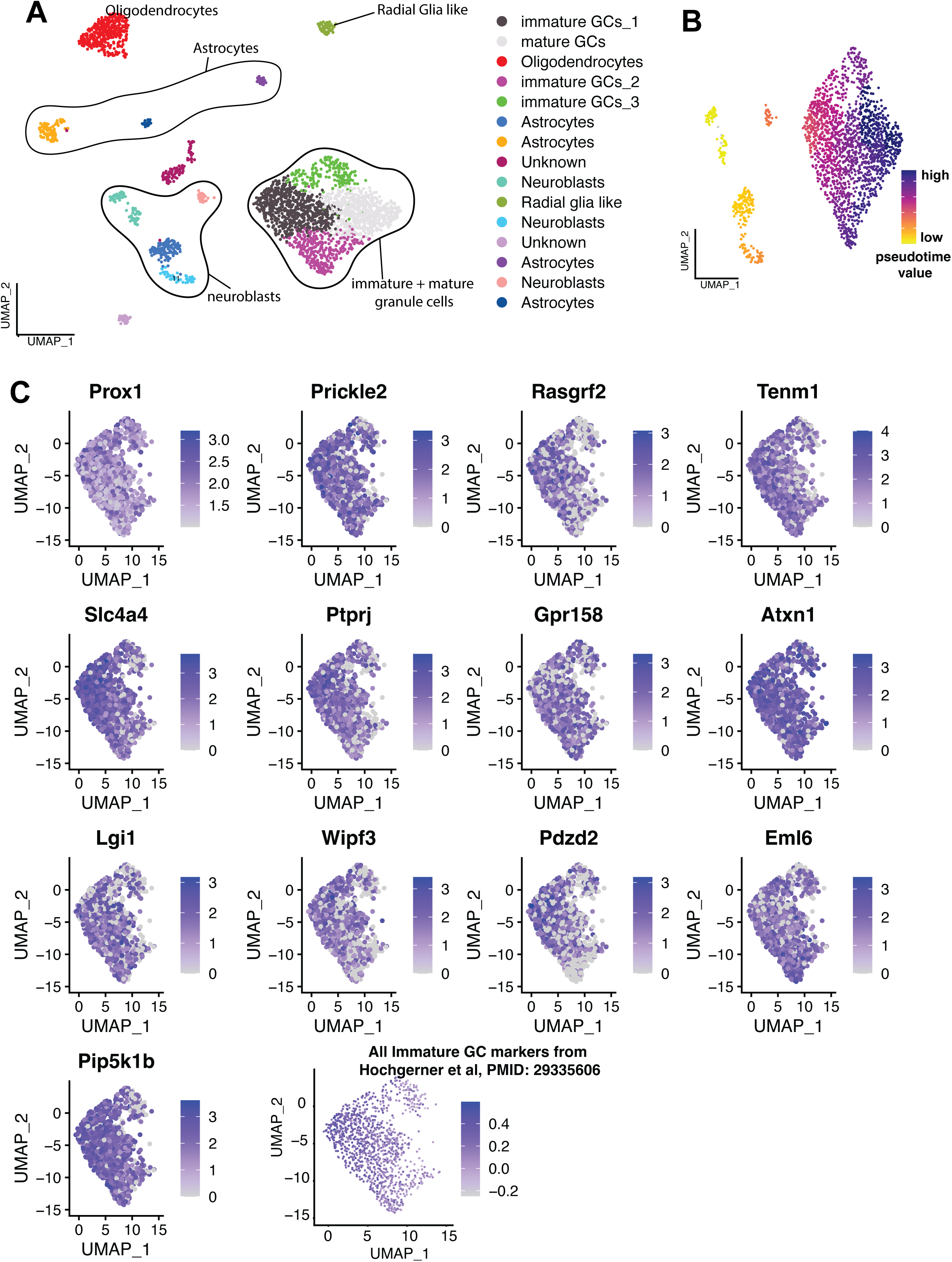
Identification of hippocampal immature GCs. A) Prox1+ hippocampal nuclei were sorted *in silico*, clustered, and visualized on a UMAP. Cell clusters were annotated based on marker genes. Neuroblasts and GCs were selected for downstream analyses. B) Pseudotime analysis was performed to gain better insights into granule cell maturation. Cells are color coded based on the inferred progression along the maturation trajectory. The color code represents the relative pseudotime values, with yellow indicating cells at the earlier stage of maturation and dark blue indicating cells at the mature stage. C) Immature GC-specific markers were used to further confirm the presence of immature GCs.

**Fig S11.**
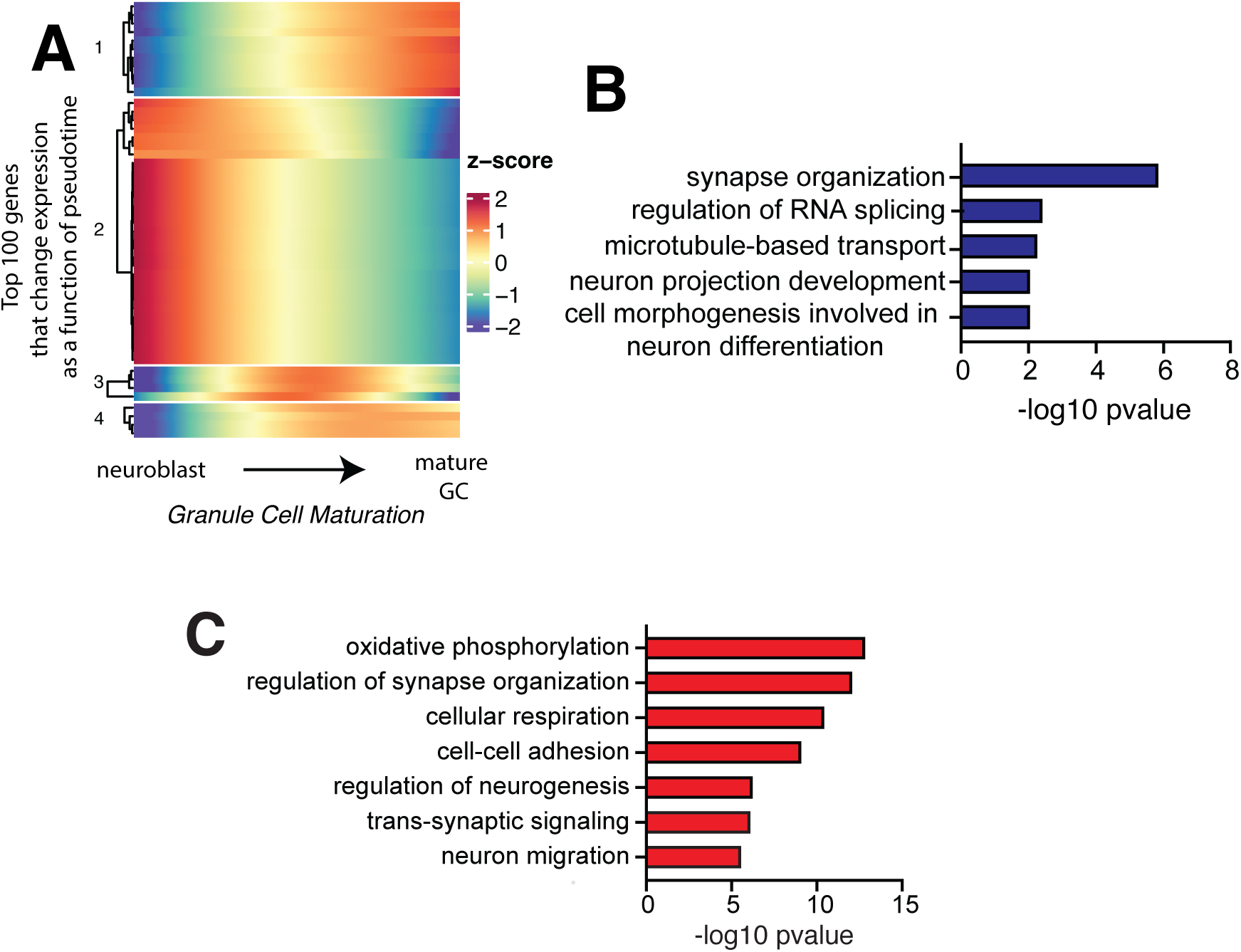
Multisensory gamma stimulation increases oxidative phosphorylation in a subset of immature granule cells. A) Heatmap showing relative expression of top 100 genes that significantly changed (q<0.05) along the maturation trajectory of granule cells (GC) in stimulation vs ambient Ts65Dn mice B) Biological processes representing differentially expressed genes (up and down-regulated) along the maturation trajectory of GC between the stimulation and ambient groups. C) Biological processes for genes differentially expressed between cluster 1,3 and cluster 2,3.

## Supporting Information

**Table S1A-B:** List of differentially expressed genes in excitatory and inhibitory neurons after stimulation.

**Table S2:** Gene enrichment analysis between upregulated genes in excitatory neurons and synaptic genes.

**Table S3A-B:** List of downregulated genes in Down Syndrome.

**Table S4:** List of co-expressed module genes identified based on weighted co-expression analysis.

**Table S5A-C:** List of downregulated genes in aging.

**Table S6A-D:** List of downregulated genes in Alzheimer’s Disease.

**Table S7:** Top 10 transcription factors associated with module Ex-7.

**Table S8:** List of differentially expressed following 40Hz stimulation along the granule cell developmental trajectory.

